# Using fibre-optic sensing for non-invasive, continuous dendrometry of mature tree trunks

**DOI:** 10.1101/2024.05.10.593544

**Authors:** Martijn van den Ende, Eléonore Oberlé, Thierry Améglio, Robin Ardito, Gildas Gâteblé

## Abstract

Dendrometry is the main non-invasive macroscopic technique commonly used in plant physiology and ecophysysiology studies. Over the years several types of dendrometric techniques have been developed, each with their respective strengths and drawbacks. Automatic and continuous monitoring solutions are being developed, but are still limited, particularly for non-invasive monitoring of large-diameter trunks. In this study, we propose a new type of automated dendrometer based on distributed fibre-optic sensing that continuously measures the change in stem circumference, is non-invasive, and has no upper limit on the trunk diameter on which it can be installed. We perform a three-month validation experiment during which we deploy a fibre-optic cable at three localities around the trunks of two specimens of *Brachychiton*. We verify the accuracy of this new method through comparison against a conventional point-dendrometer, and we observe a consistent time lag between the various measurement locations that varies with the meteorological conditions. Finally, we discuss the feasibility of the fibre-based dendrometer in the context of existing dendrometric techniques and practical experimental considerations.

## 1 Introduction

Dendrometry plays a prominent role in plant physiology. Current techniques that measure stem diameter comprise circumferentially-averaged and local point measurements [Drew and Downes, 2009]. Out of these methods, (non-automated) dendrometer bands, tape and calipers [Liu et al., 2011, McMahon and Parker, 2015], require manual intervention for each measurement, resulting in a less than desired temporal resolution and non-uniform time sampling which preclude the use of conventional signal processing methods (such as frequency-band filtering). Moreover, since climatic variables are non-stationary and the tree response to these can be highly nonlinear [Wilmking et al., 2020], ideal measurement systems should be long-term stable and continuous in time, in order to track the response of a specimen over time. However, automated and continuously-recording sensors require signal acquisition, conditioning, and storage or telemetry systems that are adapted to the (harsh) outdoor conditions of *in-situ* studies, a feat that is not trivial to accomplish.

Variation in branch diameter can be monitored continuously, non-invasively and in real time using continuous dendrometers [Ameglio and Cruiziat, 1992, Zweifel et al., 2000, 2006]. On a daily basis, the variation in branch diameter is the result of several components, including irreversible radial growth, reversible dehydration/rehydration of living cells, thermal expansion and contraction, and the expansion of conducting elements due to the increase and relaxation of internal tension [Daudet et al., 2005, Steppe and Lemeur, 2004, Zweifel et al., 2006]. In the event of severe drought, radial growth is halted and the variation in branch diameter mainly represents the fluctuation in water storage in elastic tissues, including most living cells. [Andriantelomanana et al., 2024, Lamacque et al., 2020, Zweifel, 2016, Zweifel et al., 2016]. Dendrometers have also given us more insight into trees’ resistance to freezing and the consequences of the freezing and thawing processes [Ameglio et al., 2001, Charra-Vaskou et al., 2016, Charrier et al., 2017, Zweifel and Häsler, 2000]. Dendrometry can be complemented with other types of instrumentation, such as stomatal conductance analysis [Hernandez-Santana et al., 2023, Marchin et al., 2022], sap flow measurements [Iqbal et al., 2021, Steppe and Lemeur, 2004], and water potential measurements [Cochard et al., 2001, Moriana et al., 2012], to provide a more holistic view of the response of the specimen.

However, as these techniques represent local point-measurements, they may yield biased measurements if the stem diameter variation is non-isotropic along the circumference of the stem. This may arise when fluid transport is not homogeneous due to predominant positioning of branches, or when wind triggers the formation of flexure wood and ovalisation of the stem [Roignant et al., 2018]. Another case is given by the non-uniform mechanical response of the bark, e.g. in species with a thick, scaley rhytidome (dead layers of old periderm), commonly seen in pines, or with the outer bark shedding off in sheets (Platanus L. and Eucalyptus L’Hér.) [Coder, 2022]. In these and many other cases, it is desirable to measure the stem diameter variation averaged over the entire circumference, which may be a more representative measure for comparison between specimens within or between species.

Moreover, it is often impractical (or even impossible) to install instrumentation along large-diameter stems. Around the world, there exist many veteran tree specimens that have a particular historical, cultural, or botanical interest. Local, national, and international initiatives intended to preserve this biological heritage (such as the *arbres remarquables* in France, Great trees or Champion Trees in the US and UK [Roesch et al., 2024], and heritage trees in China [Jim, 2017]) have a need for affordable and reliable monitoring of the health of the specimen, as well as its response to climatic changes, resulting in e.g. extended hydraulic stress. The use of invasive and destructive techniques on these specimens is avoided, particularly for specimens that are fragile. To some extent, automated band dendrometers alleviate many of the aforementioned issues, though their accuracy has been found to be rather limited due to friction, hysteresis, and the difficulty of eliminating slack during installation [Herrmann et al., 2016, Wang and Sammis, 2008]. Hence, there is a clear niche for a new type of dendrometric instrumentation that can be applied in the scenarios described above.

Relatively recently, distributed fibre-optic sensing [Hartog, 2017] has emerged as a promising technology for measuring strain (and temperature) in a distributed manner. By systematically sending short pulses of laser light into an optical fibre (as commonly used in telecommunication), local strain can be inferred as a function of distance along the fibre through interferometric or spectral analysis techniques. Various sensing techniques have already been adopted in a range of scientific disciplines, including engineering [Hubbard et al., 2021, Liu et al., 2023], geophysics [Jousset et al., 2018, Lindsey and Martin, 2021, van den Ende and Ampuero, 2021], and ocean monitoring [Bouffaut et al., 2022, Rivet et al., 2021], and to a limited extent, in ecology [Ashry et al., 2020]. Fibre-optic cables are flexible, robust to environmental factors like temperature and UV radiation, and are relatively inexpensive. These characteristics unlock a wide range of applications and targets for deployment.

Recognising the potential of fibre-optic sensing applied to dendrometry, we propose a novel experimental set-up to continuously record the stem diameter variations of tree specimens. More specifically, we conducted a 3-months long experiment measuring the radial expansion of two specimens of the genus *Brachychiton* with Brillouin-based Distributed Strain and Temperature Sensing (DSTS). In what follows, we describe in detail the DSTS methodology, the experimental configuration, and the data processing workflow. We interpret our observations of trunk strain variations in the context of the meteorological conditions encountered during the experiment, and the known differences in responses to hydraulic stress between the two *Brachychiton* species [Reynolds et al., 2018]. We conclude with an assessment of the feasibility of Brillouin-based DSTS for distributed dendrometry and with perspectives for the future.

## 2 Methodology

### 2.1 Description of study site and species

The data were collected from 28 July to 02 November 2023 at the Villa Thuret Botanical Gardens in Antibes, South-East France (43°33’47.8”N, 7°7’30.6”E, 40 m a.s.l). The garden was established in 1857 by Gustave Thuret, and is now managed by INRAE (France’s National Research Institute for Agriculture, Food and Environment), a public research institute. It spans an area of 3.5 ha and contains a collection of 2500 plant specimens, representing 1000 species and 131 families. Pruning is only done for safety reasons and acclimation to the local climate is tested by restricting artificial watering to the first three years after planting [Ducatillion and Landy, 2010]. Hence, the specimens present on this site are adapted to the local, coastal Mediterranean climate [Csa Köppen subtype; Kottek et al., 2006]. This climate is mainly defined by drought in summer, mild winters, and precipitations concentrated in autumn and spring [Daget, 1977]. The typical rainfall in Antibes averages 800 mm per year, although it only reached 530 mm in 2022. The average annual temperature is 15 °C, with a minimum and maximum around 0 °C and 34 °C, respectively. Locally, the clay-rich soil layer extends only to shallow depths (60 cm to 80 cm), and owes its elevated pH (*>* 7) to an important quantity of calcium [Gras, 1990].

The specimens studied in this work are from the genus *Brachychiton* Schott & Endl. (Malvaceae family), which are known to tolerate drought and to grow in varied climates and soil types [Elliot and Jones, 1985]. The first, *Brachychiton discolor* F.Muell. (Lacebark tree), has a typical height of 10 m to 30 m, is deciduous before flowering, and grows in rain forests and scrubs of coastal eastern Australia (Qld, NSW). The second, *Brachychiton populneus* (Schott & Endl.) R.Br. (Kurrajong tree), has a typical height of 6 m to 20 m, is evergreen but can be drought deciduous, and is found in rocky areas, riverbanks and semi-arid plains of inland SE Australia (Qld, NSW, Vic) [Elliot and Jones, 1985]. *Brachychiton populneus* is considered to be slightly more drought tolerant than *B. discolor* [Reynolds et al., 2018]. Both specimens tested at the Villa Thuret are mature, and their heights are 9.5 m, for *B. discolor* and 10 m for *B. populneus*. Their DBH are 136 cm and 115 cm respectively.

### 2.2 Distributed Strain and Temperature Sensing

As a subcategory of fibre-optic sensing, Distributed Strain and Temperature Sensing (DSTS, also known as Brillouin Optical Time-Domain Reflectometry, B-OTDR) is ideally suited to long-term stable monitoring of both fibre strain and ambient temperature. As described in detail by e.g. Hartog [2017], the measurement principle of DSTS is based on spontaneous Brillouin scattering, which arises from the interaction between an incident photonic wave (the probe wave), and thermally-excited acoustic vibrations in the fibre lattice called phonons. Those phonons whose wavelength matches that of the probe wave interfere with the photons to produce back-scattering that can be detected by the instrument at one end of the fibre. The Brillouin back-scattering frequency spectrum exhibits a peak with intensity *I* at frequency *ν*, which relate to fibre strain *ε* and temperature *T* as:

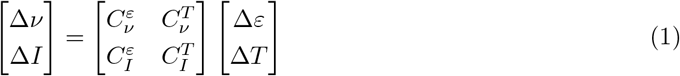

By measuring the peak frequency and intensity relative to an arbitrarily defined baseline, Δ*ν* and Δ*I* can be converted into strain and temperature relative to that same baseline (Δ*ε* and Δ*T*, respectively). Typical values for the coefficient matrix have been established by Maughan et al. [2001] as:

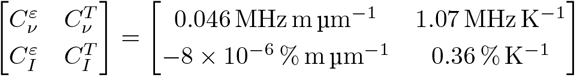

For a given relative spectral measurement (Δ*ν*, Δ*T* ), the physical quantities (Δ*ε*, Δ*T* ) are obtained by inverting this (constant) coefficient matrix and left-multiplying. As we have not calibrated these coefficients to account for the specific thermo-mechanical properties of our fibre-optic cable assembly, we directly utilise Δ*ε* estimated during acquisition, and correct for any temperature dependencies through the use of reference cable segments (this will be described in detail in the next section).

A single DSTS measurement is distributed over the fibre, meaning that during a single acquisition cycle, a Brillouin spectrum is obtained at discrete points that are evenly distributed in space (the spacing of which is referred to as the gauge length *L*). Optionally, the gauge length can be oversampled to obtain more sensing points along the fibre, although this practice does not directly lead to a higher spatial resolution as two sensing points within one gauge length are statistically dependent. Hence, DSTS provides measurements of Δ*ε* that are averaged over a gauge length. By repeating the acquisition cycle every Δ*t* time units, a data set of (relative) strain is obtained with a spatial resolution *L* and a temporal resolution Δ*t*.

### 2.3 Experimental set-up

We performed a small-scale experiment by deploying 100 m of tight-buffered single-mode fibre (type: G652.D) around the trunk of two trees; first, several winds of fibre were tightly looped around the trunk of a *Brachychiton populneus* (location B in Fig. 1, at a height of 85cm), followed by several more tight winds one metre higher up the same trunk (location A in Fig. 1, at a height of 185cm). Since DSTS is simultaneously sensitive to strain and temperature, additional fibre was wound very loosely around location A. Owing to the negligible coupling between this loose fibre and the trunk, this section of fibre is sensitive only to temperature, and can hence be used to correct the measurements of the tightly-wound sections. While Eq. (1) should (in theory) be able to separate the effects of strain and temperature, we find that this separation is imperfect and that in practice the use of a reference section is preferred. Finally, the fibre was looped (both tightly and loosely) around a second tree (*Brachychiton discolor* ; location C in Fig. 1, at a height of 160cm), which was equipped with an independent LVDT-based micro-dendrometer (described in more detail below).

**Figure 1.**
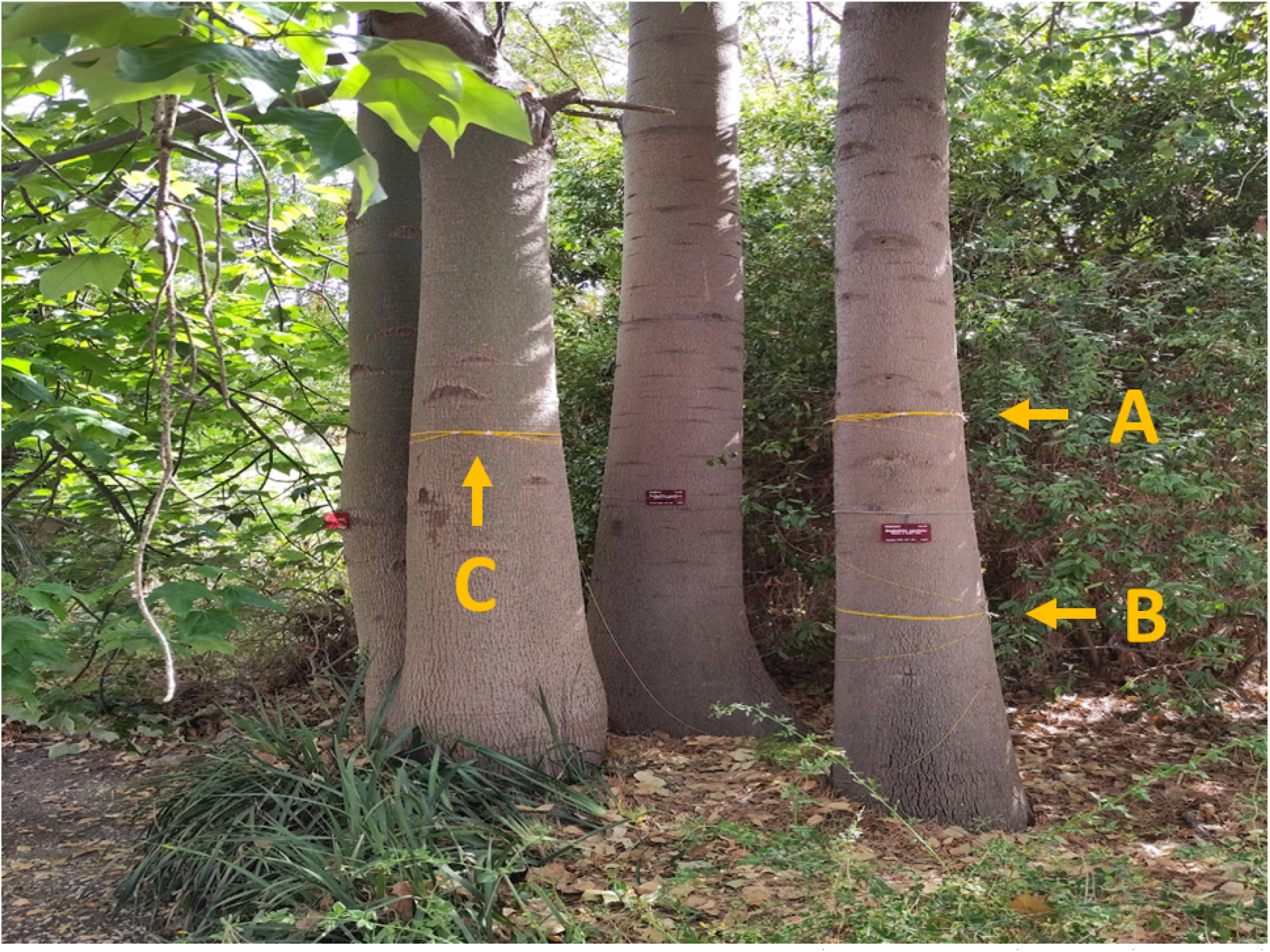
Overview of the fibre deployment. Fibre sections A (tight and loose) and B (only tight) are looped around a *Brachychiton populneus*. Fibre section C (tight and loose) is looped around a *Brachychiton discolor*, which is equipped with a PépiPIAF dendrometer installed on a branch (not in view of this picture).

With the fibre in place around the trees of study, the fibre was rolled back to starting point and connected to a Febus G1-C-19002 DSTS interrogator [Clément et al., 2021]. The section of fibre spanning between the trees and the interrogator was placed within a rubber tube for protection. The acquisition was launched with a time sampling period of 300 s and spatial resolution (gauge length) of 2 m, oversampled to obtain a spacing of 0.5 m. To geolocate specific sections of interest (rubber tube entry and exit points, tight and loose sections), tap tests were performed by manually straining the fibre at specified times, which could later be easily identified in the processed data.

As a reference instrument, we employ the PépiPIAF dendrometry system (Capt-Connect, Clermont-Fd, France), which features a linear variable differential transformer (LVDT) that continuously monitors micro-metric variations of the diameter of branches or trunks. This instrument has previously been used in various studies [e.g., Charrier et al., 2017, Daudet et al., 2005, Lamacque et al., 2020]. The specific model used in the present study is the e-Pépipiaf 1.0, which is equipped with a Solartron DF5.0 LVDT sensor, with a resolution of 0.5 μm, as well as a thermistor housed inside a waterproof case to simultaneously record ambient temperature [Chevallier et al., 2011, Lamacque et al., 2022]. The instrument is mounted on (*B. discolor* on a north-east facing branch with a diameter of 20 cm at a height of 3.25 m above the ground.

After data retrieval from the instruments, the fibre segments corresponding with locations A, B, and C were extracted. The channel recordings within a given section were averaged to reduce uncorrelated noise, and the recordings of the loosely-coupled sections (which contain the apparent strain signal due to temperature changes) were subtracted from those of the tightly-coupled sections. The resulting time-series for locations A, B, and C were thus unaffected by daily temperature fluctuations. For reference, meteorological data were obtained from the Météo-France public database for a weather station at the airport of Nice (located 10 km from the study site).

## 3 Results

Since DSTS is not yet a proven technology for dendrometry, we first consider the amplitude and long-term evolution of the radial strain of *B. discolor*. This specimen is equipped with a PépiPIAF dendrometer that has been successfully used in previous studies, and which will act as a reference sensor. Even though section C of the optical fibre was not deployed directly adjacent to the PépiPIAF, it is located sufficiently close to warrant a direct comparison. To facilitate this, the output of the PépiPIAF LVDT sensor was converted from units of millimetres to units of microstrain through division by the diameter of the branch it was mounted on (20 cm). The (equivalent) strain relative to the start of the experiment is given for the PépiPIAF dendrometer and section C of the optical fibre in Fig. 2b. While small deviations between the two time series can be occasionally discerned, overall the agreement for both the diurnal variations and the long-term evolution is more than satisfactory. Moreover, since the two sensors are not directly collocated, it is not expected to obtain perfectly identical results, and so we attribute minor amplitude discrepancies to the different response of the trunk and the branch at which the measurements are made. Particularly for the period after 21 October, corresponding to a period of net secondary growth, the trunk appears to be accumulating more strain than the branch.

**Figure 2.**
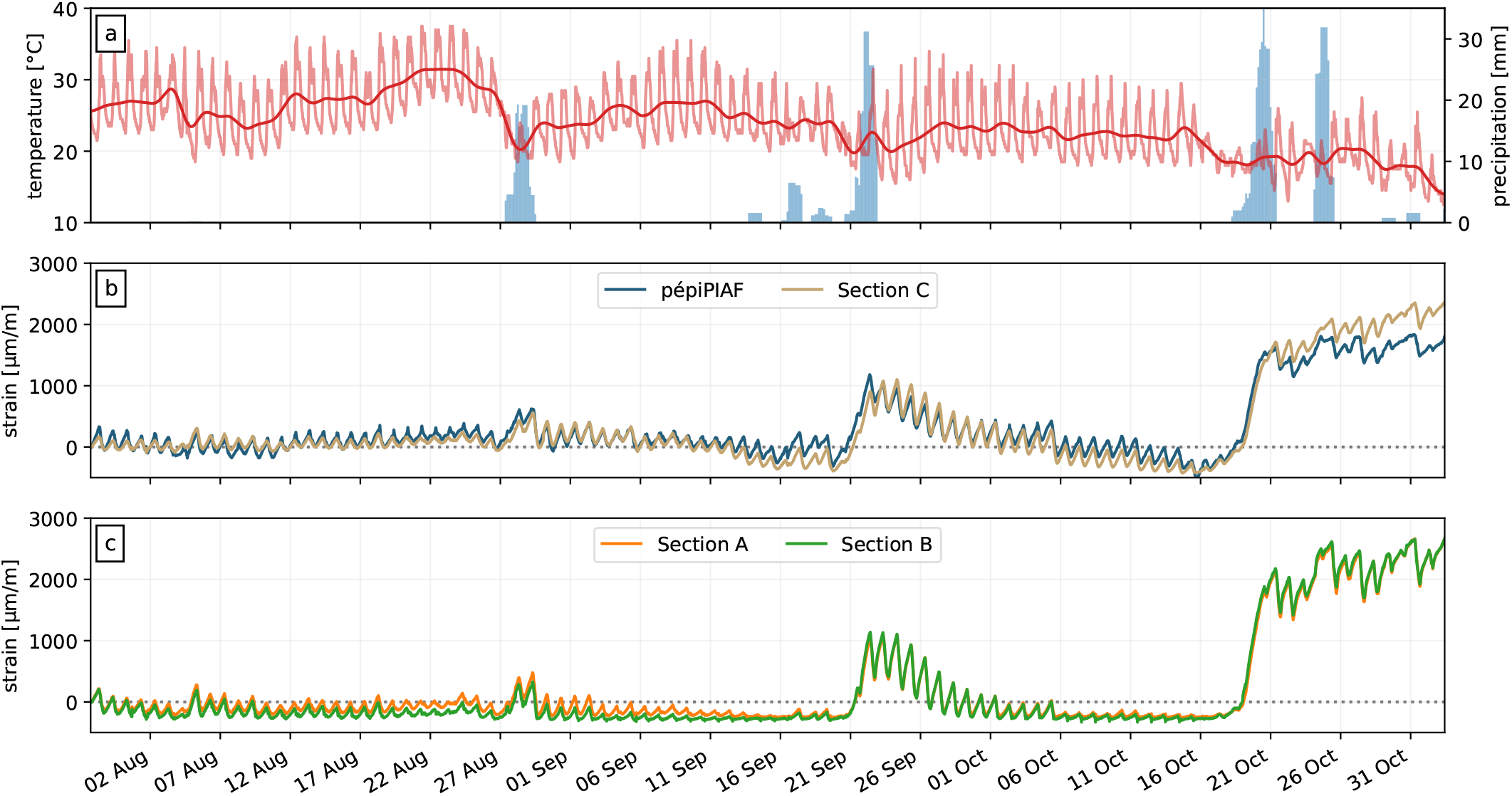
(a) Temperature recorded by the PépiPIAF on-board thermocouple and daily precipitation recorded by a weather station located ∼10 km away. (b) Strain of the fibre located at section C, and that recorded by the PépiPIAF dendrometer on a branch on the same tree (*B. discolor* ). (c) Strain of the fibre located at sections A and B (*B. populneus*). All strain measurements are relative to the start of the time series, and the dates are given in local time (UTC+2).

During the experiments, several brief episodes with a significant amount of precipitation occurred (indicated by the blue bars in Figs. 2-4). During the first episode starting on 2023-08-27 a modest amount of precipitation was measured (up to 40 mm over 48 hours), and a minor response of both *Brachychiton* specimens was recorded. The subsequent precipitation episodes were more significant, triggering a more considerable response in both specimens, dilating by as much as 1000 μm m^*−*1^ (equivalent to a radial dilatancy of the trunk of approximately 0.25 mm).

Considering the radial strain variations in more detail, we find subtle longitudinal variations in each specimen. First, when we overlay the strain recordings of the pépiPIAF dendrometer and fibre section C, we observe a small but persistent time lag of the PépiPIAF with respect to the fibre (Fig. 3a). This time lag can be effectively quantified by converting two narrowband-filtered time series into their analytic representation and subsequently extracting the phase as the complex argument. Through this analysis, we obtain the instantaneous time lag Δ*τ* (*t*) between the pépiPIAF and the fibre recordings, as well as the instantaneous amplitude *A*(*t*) of the strain signal, which we can track over time. We provide a detailed description of the analytic signal approach in Appendix I.

**Figure 3.**
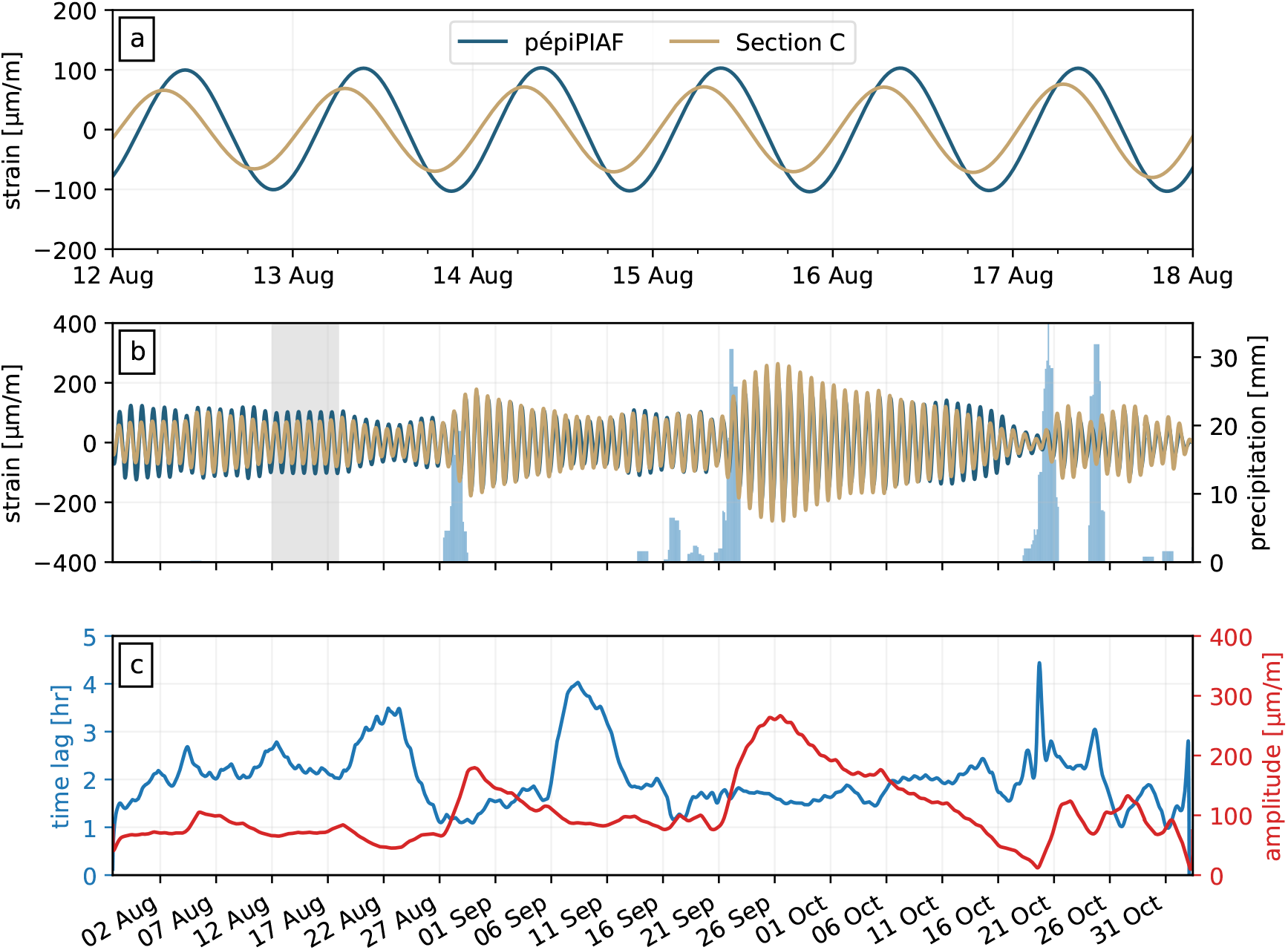
Comparison between the PépiPIAF dendrometer and fibre section C (*B. discolor* ). (a) Detailed view of the bandpass-filtered strain data, clearly showing the time delay between section C and the pépiPIAF. (b) Complete time series of the bandpass-filtered strain data. For reference, the time range of panel (a) is indicated by the shaded area, and the daily precipitation is given by the blue bars. The time series have the same colour coding as in panel (a). (c) The instantaneous time delay and amplitude of the strain.

Applying this processing procedure to the PépiPIAF and fibre recordings, we estimate the time delay to fall in the range of 1-3 hours, typically (Fig. 3c). The corresponding instantaneous amplitude of the analytic signal ranges from *<* 100 to close to 250 μm m^*−*1^. Note that owing to the narrowband filtering applied (see Appendix I), this amplitude only reflects the strain associated with the diurnal hydration cycle. When compared to the recordings of daily precipitation, a persistent increase of the time lag seems to coincide with a period of drought. More strikingly, the amplitude of the diurnal hydration cycles correlates directly with the onset of precipitation.

Performing a similar analysis on fibre sections A and B (*B. populneus*), we observe a time lag that is systematically much smaller than that seen for *B. discolor*, usually remaining below 1 hour (Fig. 4). This is not necessarily unexpected, as fibre sections A and B are both located on the trunk, separated by a distance of only 1 m, which are markedly different from the locations and distance between fibre section C and the PépiPIAF dendrometer on the other specimen. Moreover, the same trends are observed as for *B. discolor* : the time lag tends to increase during periods of drought, and the instantaneous amplitude increases dramatically with each precipitation episode.

**Figure 4.**
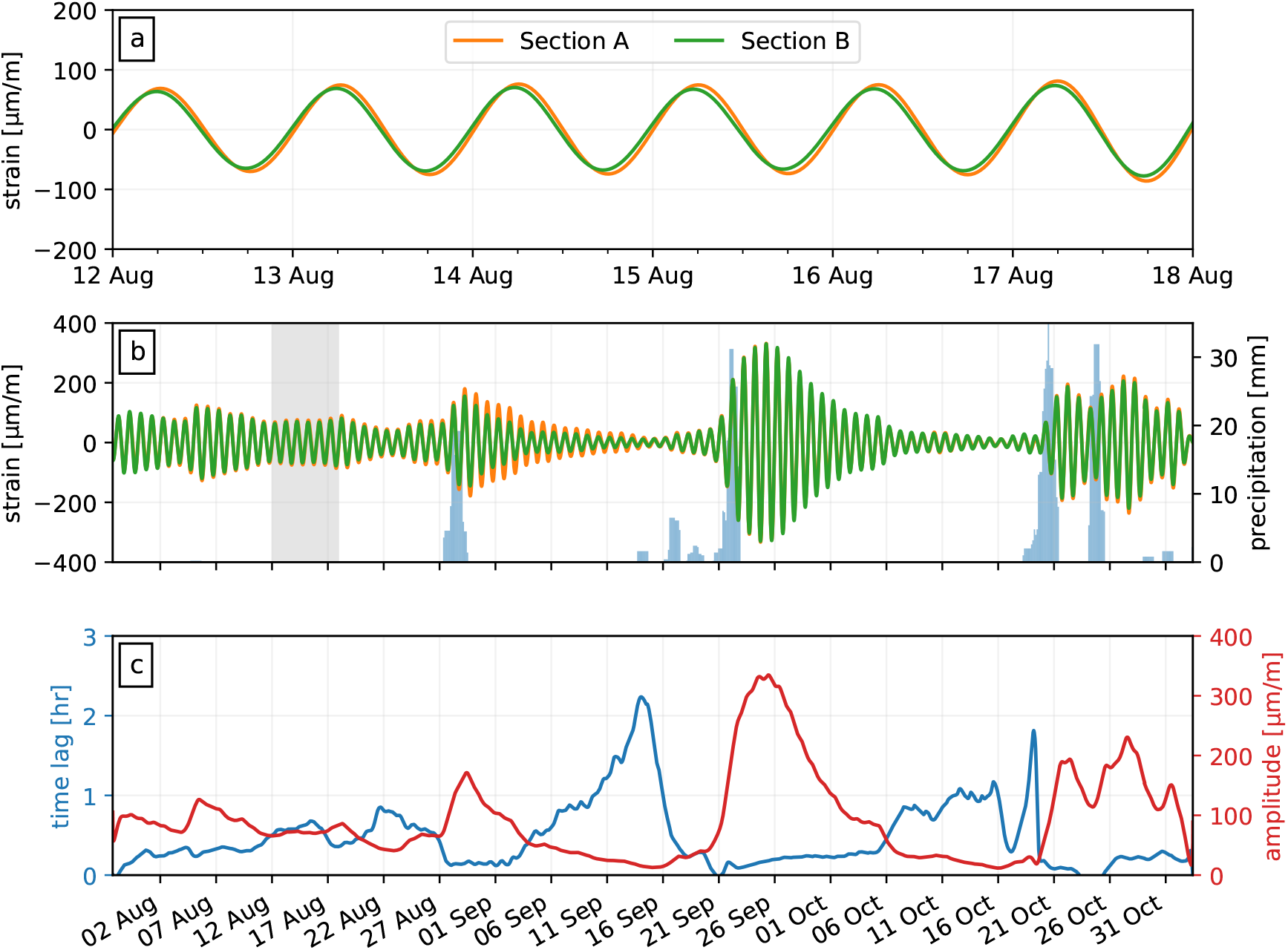
Comparison between the fibre sections A and B (*B. populneus*). (a) Detailed view of the bandpass-filtered strain data, showing a minute time delay between sections A and B. (b) Complete time series of the bandpass-filtered strain data. For reference, the time range of panel (a) is indicated by the shaded area, and the daily precipitation is given by the blue bars. (c) The instantaneous time delay and amplitude of the strain.

## 4 Discussion

### 4.1 Interpretation of hydraulic response

It is clear from our analysis that there is a direct correlation between the amplitude of the diurnal hydration cycles and the availability of water. This has been observed in previous studies [Daudet et al., 2005, Fernández and Cuevas, 2010, Steppe and Lemeur, 2004], and it is well understood in terms of an increased sap flow throughout the day [Zhao and Liu, 2010]. What can be seen particularly well in Fig. 2c, is that a larger daily amplitude corresponds to an increase of the maximum strain seen throughout a day, while the baseline diameter remains relatively constant over time, indicating a lack of secondary growth. This indicates that the amplitude variations are due to an increased use of water reserves in bark cells compared to a relatively constant storativity baseline. This constant baseline may result from drought induced stomatal closure (corresponding with minimal diurnal strain variations), in order to stabilise the water potential and avoid cavitation of the sap [Choat et al., 2018]. A maximum of fluid storage, which marks the end of the periods of rehydration and growth, is typically achieved around 08:00 local time.

In contrast to the amplitudes of diurnal dynamics, the time lag between discrete locations along the trunk is less straightforward to interpret. For both specimens we observe a systematic positive time lag, meaning that the sensor higher up on the tree (the PépiPIAF sensor for *B. discolor*, or fibre section A for *B. populneus*) exhibits a delay in the signal. Previous studies that simultaneously measured the rate of sap flow at the base of the trunk and in the crown of a specimen report that sap flow initiates near the crown, triggered by the onset of transpiration [Phillips et al., 1999, Zweifel et al., 2001]. The observed time lags of the initiation of sap flow range from hours [Čermák et al., 2007] to minutes [Kume et al., 2008]. Hence, when considering the rate of sap flow, one would expect to make the opposite observation (i.e., a negative time lag) from what is measured in the present study. However, one needs to keep in mind that with both the PépiPIAF dendrometer and with the fibre, one records radial strain, which is not necessarily directly linked with the rate of sap flow; instead, it is the net storage of fluid in the elastic sapwood and phloem tissue that contributes directly to stem diameter variations [Čermák et al., 2007]. An accumulation of sugar higher up the tree could possibly induce an osmotic pressure gradient that stimulates fluid flow towards these regions, causing storage at higher positions and depletion at lower positions on the tree. Nighttime transport of solutes towards the base of the tree, along with rehydration, would reverse this trend, hence resulting in a daily stem variation that occurs earlier in the day for lower positions. Compared with other dendrometric studies (as opposed to sap flow studies), positive or zero time lags in stem diameter or between sap flow and transpiration, have been occasionally reported [Ameglio and Cruiziat, 1992, Simonneau et al., 1993, Steppe and Lemeur, 2004, Ueda et al., 1996]. Exactly what controls the retention and release of fluids is subject of debate [Sevanto et al., 2002], and is likely highly dependent on the species under investigation [Scholz et al., 2008]. We therefore cannot be conclusive about the potential role of solutes in explaining the observed lag times.

A second candidate hypothesis is the difference in exposure to solar radiation: east facing branches (which are illuminated earlier in the day) tend to have an earlier peak in sap flow, compared to west facing and lower branches [Burgess and Dawson, 2008, Martin et al., 2001], and southern branches can have a higher total/overall sap flow than northern branches [Steinberg et al., 1990]. In this study, the PépiPIAF was mounted on a lower branch oriented to the North-East, usually in the shade of a building. Since the fibre at section C measures the stem variations averaged over the circumference of the trunk, one might anticipate time shifts when comparing a specific branch to the tree as a whole. However, this does not explain the time delay observed between sections A and B, which are both situated on the same trunk.

Regardless of the relation between sap flow rates and fluid storage, it is interesting to note that as soon as the water availability increases (during or shortly after precipitation), the observed time lag diminishes; particularly in the case of *B. populneus* this time lag approaches zero, implying that the trunk essentially acts as a rigid fluid reservoir that uniformly stores and releases fluids from and to other parts of the specimen. In between precipitation episodes, the time lag becomes non-zero, resulting in a distributed storage and release of fluids starting at the base of the trunk and propagating upward. By having several continuous measurements distributed over the length of the stem, it has become feasible to study this transition from uniform to heterogeneous fluid release in detail, particularly in relation to the availability of water and the state of hydraulic stress. In turn, such observations can be interpreted in terms of species-dependent strategies of mitigating hydraulic stress and their resilience to episodic drought [Chirino et al., 2011].

Both species studied here are known to be drought resilient, with a relatively high basal increment given the low transpiration measured. The natural range of *B. discolor* is slightly more mesic than that of *B. populneus* [McCarthy et al., 2011], which might impact their respective hydraulic stress mitigation strategies. Seedlings of *B. populneus* were found to have higher stomatal conductance than *B. discolor* in the control setting and lower in the drought setting [McCarthy et al., 2011]. This plasticity makes these seedlings better adapted to seasonal rainfall and enhances their water use efficiency during droughts [Reynolds et al., 2018]. In the present study, *B. populneus* experiences a more intense response (in terms of daily amplitude variations) to precipitation than *B. discolor*, which suggests higher evapotranspiration and growth. Additionally, after several days without precipitation, *B. populneus* exhibits a daily stem variation that approaches zero, potentially implying at least partial stomatal closure. These results are in line with previous observations made on seedlings, and support the current view on the drought resistance of these two species.

### 4.2 Feasibility of DSTS and perspectives

The favourable comparison between the DSTS strain measurements and those obtained from an established instrument (the PépiPIAF dendrometer; see Fig. 2b) demonstrates that the fibre-based strain measurements reliably record the radial expansion and contraction, over time scales ranging from hours to months. Aside from the stability and accuracy of the measurement, the signal-to-noise ratio is comparable to that of the PépiPIAF LVDT sensor, facilitating detailed analyses like those presented here. Hence, DSTS offers a viable alternative to conventional dendrometric instruments, while holding the following advantages:

- A single instrument (the interrogator) is capable of continuously sensing up to 100 km of fibre at a metric resolution, permitting the simultaneous monitoring of numerous positions on one or multiple specimens. Compared to installing, maintaining, telemetering, and synchronising the data of multiple instruments, the logistics of fibre-based instrumentation are greatly simplified.
- Fibre-optic cables are flexible yet robust to tension, resistant to heat, water, and sun exposure, and they are electromagnetically inert (since no electric current runs through them). This makes optical fibres highly suitable for long-term deployments in exposed (outdoor) settings. Moreover, with a nominal cost of the order of 1 €/$/£ per metre, the cost of replacement is minimal, and damaged sections can be readily repaired on-site. Compared to conventional electronics, fibre-optic cables require minimal protective measures for deployment in harsh conditions.
- The inherently distributed nature of DSTS enables the investigation of the hydrological dynamics in high spatio-temporal resolution. As aforementioned, follow-up studies that utilise fibre segments located at a number of localities along the stem of a tree offer an opportunity to elucidate fine-scale details of the storage and release of fluids. Moreover, the technique is completely non-invasive, and does not require human intervention beyond the initial deployment.

We therefore envision that DSTS will facilitate future long-term dendrometric monitoring programmes. Complemented with e.g. sap flow rate and hydraulic potential measurements, distributed dendrometry will lead to renewed insights into the species-specific response to climatic stressors, mitigation strategies, and long-term adaptability.

## 5 Conclusions

In this study we propose *Distributed Strain & Temperature Sensing* (DSTS) as a novel technology for long-term, continuous dendrometry. To test the feasibility of DSTS for this purpose, we deployed an optic fibre around the stems of two specimens of *Brachychiton* (*B. discolor* and *B. populneus*). *Brachychiton discolor* was equipped with a conventional dendrometric instrument (PépiPIAF sensor) that acted as a reference. From the comparison of the continuous DSTS strain recordings with those derived from the conventional dendrometer, we conclude that the DSTS strain measurements are both accurate and long-term stable, validating DSTS as a feasible methodology. Furthermore, the spatially distributed DSTS measurements reveal intriguing lateral variations in the diurnal diametric fluctuations of the stems, which we interpret to relate to the local storage and release of fluids, and variations thereof driven by the availability of water. Hence, DSTS-based dendrometry has great potential for complementing existing dendrometric techniques, and to deliver new insights into the spatially-variable hydrological response of tree species.

## Appendix I: analytic signal processing

Consider a time-domain signal *x*(*t*). The corresponding *analytic signal* 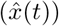 is obtained as 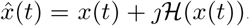, with *ℋ*(·) denoting the Hilbert transform and *ȷ*^2^ ≡ *−*1 being the imaginary unit. This analytic representation is complex-valued, and so it can also be represented in polar form as:

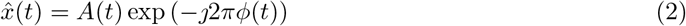

with instantaneous amplitude *A*(*t*) and phase *ϕ*(*t*). Consider now two time series, 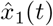 and 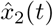 that are filtered in a narrow frequency band around central frequency *f*_0_. Multiplying 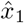 with 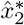 (* denoting the complex conjugate), then gives:

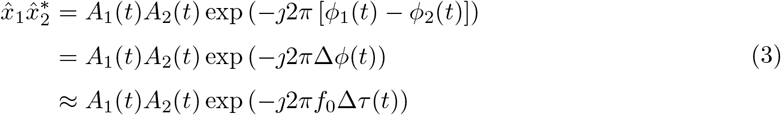

Here, Δ*τ* (*t*) denotes the instantaneous time lag between signals *x*_1_(*t*) and *x*_2_(*t*). Hence, the time lag is explicitly computed as:

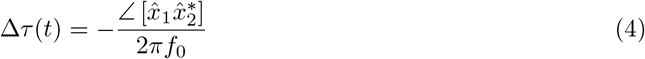

with ∠ [·] denoting the complex argument. Likewise, the instantaneous amplitude of *x*(*t*) is simply:

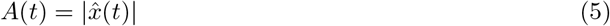

For the analysis of the diurnal hydration cycles, we set *f*_0_ corresponding to a period of 24 hours. We then apply to the strain data a non-causal peak filter centred on *f*_0_ with a dimensionless quality factor of 3, resulting in a time series that only contains the diurnal variations. We subsequently transform the filtered data into its analytic form with Eq. (2), and estimate the time lag and instantaneous amplitude with Eqs. (4) and (5), respectively. Note that the time lag inferred in this manner is independent of the time resolution of the measurement (5 minutes), and so time lags smaller than this time resolution can be correctly resolved.

## Acknowledgments

MvdE thanks A. Sladen (CNRS Géoazur) for making the DSTS instrument available for this experiment, and P. Clement (Febus Optics) for in-depth discussion of the DSTS measurement principle. EO, RA and GG thank the city of Antibes and especially P. Dalmasso for their financial support (n° C10870) to this study. The authors thank P.-E. Lauri for his feedback on the initial draft of the manuscript. For the analysis we made use of the following Python libraries: Matplotlib 3.7.2 [Hunter, 2007], NumPy 1.25.1 [Harris et al., 2020], Pandas 2.0.3 [Pandas Development Team, 2023], SciPy 1.11.1 [Virtanen et al., 2020]. All the data and scripts needed to reproduce the figures in this study are available here: https://doi.org/10.6084/m9.figshare.25773432.

